# Self-regulation of the brain’s right frontal Beta rhythm using a brain-computer interface

**DOI:** 10.1101/2021.10.07.463487

**Authors:** Nadja Enz, Jemima Schmidt, Kate Nolan, Matthew Mitchell, Sandra Alvarez Gomez, Miryam Alkayyali, Pierce Cambay, Magdalena Gippert, Robert Whelan, Kathy L. Ruddy

## Abstract

Neural oscillations, or brain rhythms, fluctuate in a manner reflecting ongoing behavior. Whether these fluctuations are instrumental or epiphenomenal to the behavior remains elusive. Attempts to experimentally manipulate neural oscillations exogenously using non-invasive brain stimulation have shown some promise, but difficulty with tailoring stimulation parameters to individuals has hindered progress in this field. We demonstrate here using electroencephalography (EEG) neurofeedback in a brain-computer interface that human participants (n=44) learned over multiple sessions across a 6-day period to self-regulate their Beta rhythm (13-20 Hz) over the right inferior frontal cortex (rIFC). The modulation was evident only during neurofeedback task performance but did not lead to offline alteration of Beta rhythm characteristics at rest, nor to changes in subsequent cognitive behavior. Likewise, a control group (n=38) who underwent training of the Alpha rhythm (8-12 Hz) did not exhibit behavioral changes. Although the right frontal Beta rhythm has been repeatedly implicated as a key component of the brain’s inhibitory control system, the present data suggest that its manipulation offline prior to cognitive task performance does not result in behavioral change. Thus, this form of neurofeedback training of the tonic Beta rhythm would not serve as a useful therapeutic target for disorders with dysfunctional inhibitory control as their basis.

## 1 Introduction

The synchronous firing of large populations of neurons across distributed brain networks produces rhythmic electric field fluctuations large enough to be detected at the scalp. The role – either causal or epiphenomenal – of the observed neural oscillations (‘brain rhythms’) for human behavior has been a topic of intense debate for decades. Traditionally, researchers have recorded neural oscillations from the scalp while participants perform cognitive tasks, thus investigating the correlation between brain signals and behavior. However, experimental manipulation is necessary in order to specify a causal role for neural oscillations (Herrmann et al., 2016; Vosskuhl et al., 2018).

Exogenous modulation of neural oscillations has previously been achieved using non-invasive brain stimulation techniques like transcranial alternating current stimulation (tACS, see Vosskuhl et al., 2018 for a review), oscillatory transcranial direct current stimulation (o-tDCS; e.g., Marshall et al., 2006), and repetitive transcranial magnetic stimulation (rTMS, e.g., Chung et al., 2015; Thut & Miniussi, 2009) (for reviews, see Dayan et al., 2013; Thut et al., 2011). These methods have generated mixed results with regards to effectiveness of neural modulation (for reviews, see Demirtas-Tatlidede et al., 2013; Enriquez-Geppert et al., 2013; Polanía et al., 2018; Thut et al., 2011) and impact upon cognitive behavior (Bestmann et al., 2015) with a key issue being heterogenous responses to the same stimulation across individuals (Adeyemo et al., 2012; Bergmann & Hartwigsen, 2020; Kasten et al., 2019). Individuals exhibit subtle idiosyncratic features of brain rhythms even within the commonly described bandwidths (Benwell et al., 2019; Haegens et al., 2014), however, methods like rTMS, tACS or o-tDCS typically target specific frequencies.

In addition to neuromodulation methods, brain-computer interface (BCI)-based neurofeedback can be used to endogenously train volitional modulation of brain signals. This approach enables participants to self-regulate brain rhythms which are intrinsic to the individual brain (Ros et al., 2010). Neurofeedback has been frequently tested and used as a therapeutic tool and studies have shown behavioral improvements in disorders such as attention-deficit/hyperactivity disorder (ADHD; Jean Arthur Micoulaud-Franchi et al., 2014; Sonuga-Barke et al., 2013). However, to date there has been wide heterogeneity in research designs for neurofeedback protocols and it is difficult to draw conclusions regarding the effectiveness of neurofeedback to modify behavior (Omejc et al., 2019; Simon et al., 2021; Sitaram et al., 2017). As well as targeting individually tailored neural oscillation frequencies, it is important to target brain regions that are directly instrumental to the behavior under investigation. For example, electroencephalography (EEG)-neurofeedback from brain signals recorded over sensorimotor areas has shown some evidence of motor-skill improvement in healthy participants as well as clinical motor symptoms in ADHD or stroke patients (Jeunet et al., 2019). In addition, Hsueh et al. (2016) showed that neurofeedback of the frontoparietal Alpha rhythm improved working memory. Neurofeedback training of the Beta (13-20 Hz) rhythm in the past has predominantly targeted sensorimotor areas (e.g., Boulay et al., 2011; Vernon et al., 2003; Witte et al., 2013).

Inhibitory control is a core component of healthy executive function, and deficiencies with this aspect of cognition manifest in disorders such as ADHD (e.g., Lijffijt et al., 2005) or addiction (e.g., Luijten et al., 2011). Inhibitory control is believed to rely on fast and flexible command of the brain’s Beta rhythm (Enz et al., 2021; Jana et al., 2020; Schaum et al., 2021; Swann et al., 2009; Wagner et al., 2017; Wessel, 2020), primarily in a pathway connecting right inferior frontal cortex (rIFC) and basal ganglia (Aron et al., 2014; Wessel & Aron, 2017). The Stop Signal Task (SST) measures this cognitive process (Logan & Cowan, 1984) by requiring the participant to cancel an already initiated motor response following an infrequent Stop cue. The Stop Signal Reaction Time (SSRT) is an estimation of the covert latency of the action cancellation process (Verbruggen et al., 2019).

In order to test whether selective self-regulation of specific brain rhythms could modulate specific cognitive processes, we designed a protocol whereby 82 participants learned over a 6-day period to either upregulate or downregulate the amplitude of their Beta or Alpha rhythm using direct neurofeedback in a BCI. We measured two distinct aspects of cognitive function; speed of proactive response inhibition (conditional SST [cSST]) and working memory (2-Back Task). In a double-blinded mixed design with between-subject (neurofeedback types) and repeated measures (pre-post neurofeedback cognitive measures) factors, we tested the theory that causal manipulation of brain rhythms following neurofeedback training would have an observable impact upon behavior. Specifically, we hypothesized that learning to modulate the Beta rhythm over rIFC would impact positively upon speed of response inhibition, but not upon working memory. By contrast, we predicted that the control group undertaking Alpha neurofeedback would demonstrate no improvement in response inhibition.

## 2 Method

### 2.1 Participants

82 healthy adult human volunteers (age: 24.27 ± 7.74 years [mean ± SD]; 44 female; 69 right handed) participated in the study. Participants were reimbursed with either €5/hour and a completion bonus of €60 or with ECTS points. Inclusion criteria were: aged over 18, no history of traumatic brain injury and not currently experiencing any psychiatric disorder (self-reported). All participants provided written informed consent prior to participation. The experimental procedures were approved by the School of Psychology ethics committee of Trinity College Dublin and conducted in accordance with the Declaration of Helsinki.

### 2.2 Study design

Participants were allocated at random into four groups: ‘*Beta UP*’(n=26), ‘*Beta DOWN*’ (n=18), ‘*Alpha UP*’ (n=23) and ‘*Alpha DOWN*’ (n=15). ‘Beta’ groups were trained to either increase (‘UP’) or decrease (‘DOWN’) their Beta (13-20 Hz) rhythm over the rIFC whereas ‘Alpha’ groups were trained to either increase (‘UP’) or decrease (‘DOWN’) their Alpha (8-12 Hz) rhythm over the same region. Each participant was trained over six sessions (‘*S1-6*’). Where possible, sessions were scheduled for a similar time of the day.

The study design is illustrated in **Figure 1**. Each session consisted of one calibration (‘*Cal S1-6*’) and four neurofeedback training blocks (‘*B1-6.1-4*’) and each block lasted three minutes. Resting EEG was recorded in each session before (‘*Rest S1-6 Pre*’) and after (‘*Rest S1-6 Post*’) the four neurofeedback blocks. During S1 and S6, participants additionally performed two behavioral tasks, the cSST (experimental task) and the 2-Back Task (control task). The cSST was performed twice in S1, once before (‘*cSST S1 Pre*’) and once after (‘*cSST S1 Post*7#x2019;) neurofeedback training whereas the 2-Back Task was only performed once (‘*2-Back S1 Pre*’). In S6, both tasks were performed once after neurofeedback training (‘*cSST S6 Post*’, ‘*2-Back S6 Post*’). For all sessions and tasks, the participants were seated comfortably in a chair in front of a computer screen in a soundproof, darkened room.

**Figure 1:**
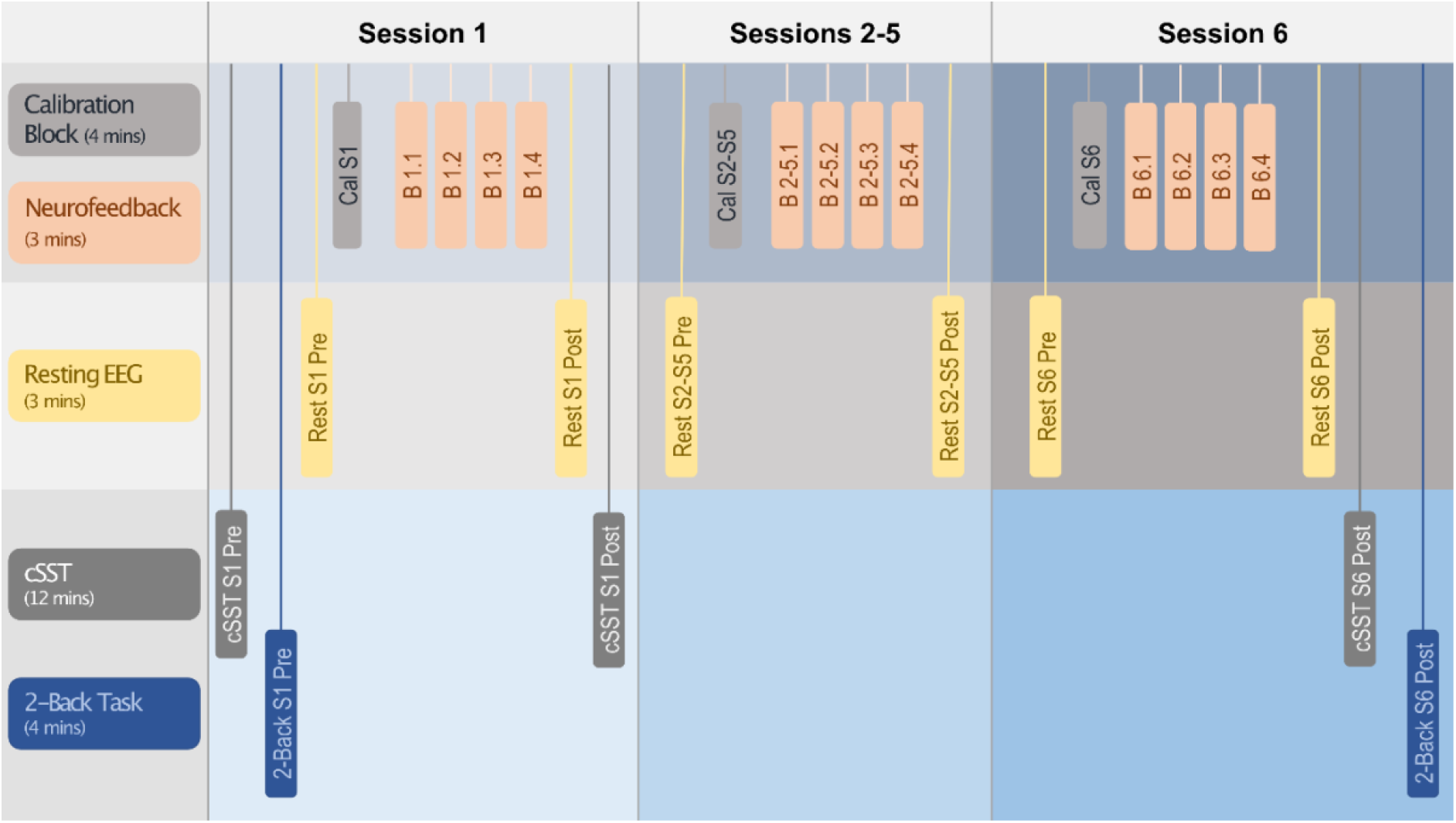
Study design. Behavioral tasks, baseline EEG measurements and neurofeedback training blocks are shown in sequential order for each of the six sessions. Each session was performed on a separate day. Abbreviations: cSST: conditional stop signal task.

#### 2.2.1 EEG recording

During S1 and S6, 128-channel EEG data in the 10-5 system format were recorded using a 128-channel BioSemi headcap connected to an ActiveTwo Biosemi system. During S2-S5, EEG was recorded from four active Ag/AgCl electrodes over the individual rIFC locus (see ‘*Functional rIFC localization*’) with the same hardware. A reference electrode recorded data from Cz. For all sessions, three additional electrodes recorded the electrooculogram (right outer canthi for horizontal eye movements and ~2 cm below the left eye for vertical eye movements) as well as the right masseter muscle to detect facial movements.

#### 2.2.2 Resting EEG

To record resting EEG, participants were instructed to fixate upon a black cross for 3 minutes with their eyes open and in a relaxed and stable body position.

#### 2.2.3 Conditional Stop Signal Task (cSST)

The cSST was performed using Presentation^®^ (Version 18.0, Neurobehavioral Systems, Inc., Berkeley, CA, www.neurobs.com) and EEG was recorded during the task. Each trial lasted 1000 ms and was preceded by a fixation cross (1000 ms duration). During Go trials, participants were presented with black arrows pointing either to the right or left (Go signal; 750 ms duration) and they were instructed to respond with their right or left index finger, respectively, as fast as possible via an Xbox 360 game controller. In one of four Go trials, the Go signal was followed by a black arrow pointing upwards (Stop signal; 250 ms duration) after a varying stop-signal delay (SSD). The participants were instructed to inhibit their button press on these Stop trials, but only if the Go signal was pointing in the critical direction. If a Stop signal appeared after a Go signal pointing in the non-critical direction, the participants were instructed to ignore the Stop signal and respond with a button press. The task was divided into four blocks; for the first two blocks the critical direction was right and for the last two blocks the critical direction was left. Each block consisted of 24 trials (18 Go trials and 6 Stop trials). The number of right and left pointing Go signals was equal in each block and presented in a randomized manner. The SSD was adjusted by a tracking algorithm, aiming to achieve a task difficulty resulting in 50% successful and 50% failed Stop trials. After a successful critical Stop trial (ignoring non-critical Stop trials), the SSD was increased, making the task harder and after a failed critical Stop trial (ignoring non-critical Stop trials), the SSD was decreased, making the task easier. The initial SSD was 250 ms and was subsequently adjusted using a double-limit algorithm (see Richards et al., 1999). The SSD could vary between 50 ms and 450 ms. Following a Stop trial, the subsequent SSD value was chosen randomly between the current SSD and a pair of limits (higher or lower, as appropriate). These limits were designed to converge on the SSD that produced a 50% success rate and to be robust to fluctuations on individual trials. If a participant responded to the Go signal before Stop signal presentation, then the SSD was decreased for subsequent trials. All participants completed one block of fifteen practice trials where they received feedback before the real task.

#### 2.2.4 2-Back Task

The 2-Back Task was programmed in Presentation^®^ and no EEG was recorded. Participants were serially presented with randomized letters (500 ms duration each) with an interstimulus interval of 1000 ms. Participants were instructed to press the keyboard button number 1 if the current letter matched the letter presented 2 letters ago. Each block contained 10 targets (i.e., matches) with the target frequency balanced across the block. The task consisted of two blocks and each block contained 50 letters.

#### 2.2.5 Functional rIFC localization

After completion of the *cSST S1 Pre*, the raw EEG data recorded during the task was subjected to an immediate analysis in order to localize the rIFC region on the participant’s topography. While scalp EEG cannot identify the source generator, studies using electrocorticography (Swann et al., 2009; Swann et al., 2012), magnetoencephalography/functional magnetic resonance imaging (fMRI; Schaum et al., 2020) and fMRI-guided repetitive transcranial magnetic stimulation (Sundby et al., 2021) have empirically demonstrated the link between right frontal scalp activity and rIFC. EEG data from the cSST were epoched into 2500 ms epochs with respect to the Go/Stop signal for Go trials and Stop trials, respectively. A short version of the Fully Automated Statistical Thresholding for EEG artefact Rejection plug-in (FASTER; Nolan et al., 2010; https://sourceforge.net/projects/faster/) was used to identify the most significant artefacts (e.g., eyeblinks and idiosyncratic muscle movements). EEG data were filtered at 13-20 Hz (Beta rhythm) and the amplitude was squared to obtain power estimates. Preprocessed EEG data were epoched into 100 ms to 300 ms after the Stop signal for successful and failed Stop trials. Additionally, a baseline (resting EEG data) was extracted from 1800 ms to 2000 ms after the Stop signal. These data were averaged over all trials for each channel and illustrated using topoplots. Four topoplots were shown; successful Stop trials, failed Stop trials, successful minus failed Stop trials and successful Stop trials minus baseline. The topoplots were then visually inspected and four right frontal electrodes showing the largest right frontal power increase in the Beta range were identified as the individual participant’s rIFC region. This four-electrode cluster was used subsequently to provide signals for the neurofeedback.

#### 2.2.6 Calibration and neurofeedback blocks (B1-4)

The OpenViBE Acquisition Server (OpenViBE, Renard et al., 2010, www.openvibe.inria.fr) received the EEG stream, and data were processed in real time using a custom OpenViBE Designer script. Data were processed using a 100-ms sliding window. First, the four individual rIFC channels as well as the reference channel (Cz) were selected. The selected data was then spatially (averaged over four rIFC channels) and temporally (13-20 Hz for ‘Beta’ groups, 8-12 Hz for ‘Alpha’ groups) filtered. The data were then epoched into 100 ms windows and a power estimate of each 100 ms epoch was calculated by squaring the amplitude. Using LabStreamingLayer (https://github.com/sccn/labstreaminglayer), the resulting power estimates were then exported to MATLAB (R2017b, Mathworks, USA). Using a custom MATLAB script, the power estimates were visualized using the Psychophysics Toolbox (PsychToolbox; Brainard, 1997; http://psychtoolbox.org).

##### Calibration

During the calibration block, the participants were instructed to first rest and fixate upon a red cross for 2 minutes and when the fixation cross turned green after 2 minutes, they were instructed to open and close their left hand for another 2 minutes. EEG was measured during the whole duration of the calibration block. This calibration procedure was performed in order to measure the participant’s full power range of the respective frequency band (Beta or Alpha). The median power of the entire block was calculated.

##### Neurofeedback training (B1-4)

During the four neurofeedback blocks, an avatar (bird for UP groups, fish for DOWN groups) was visualized on the screen and represented the participant’s real-time power estimate output from OpenViBE. The avatar moved horizontally from left to right of the screen and either upwards or downwards depending on whether the power estimates increased or decreased, respectively. The screen was separated horizontally by the median power from the calibration block into sky (above median) and sea (below median) (**Figure S2**). The top of the screen was equal to the maximum power of the calibration block and the bottom of the screen was equal to the minimum power of the calibration block. The UP groups were instructed to keep the bird in the sky (i.e., increase the power estimates) whereas the DOWN groups were instructed to keep the fish in the sea (i.e., decrease the power estimates). If the avatar deviated into the wrong environment (i.e., sky or sea), the environment turned red to give negative feedback and immediately turned back to normal when they returned into the desired zone. When participants were frequently reaching the top (UP groups) or bottom (DOWN groups) of the screen, the minimum and maximum limits were expanded by the X*standard deviation of the power estimates of the calibration block. The game thus had 4 difficulty levels (X=0-4) and the level was increased if the participant was able to stay in the correct area (sky or sea) for more than 95% in a block. The participants were instructed to develop a mental strategy that does not involve movements, clenching teeth or tensing muscles. Participants were instructed to not close their eyes and to fixate upon the screen at all times.

### 2.3 Data offline processing

#### 2.3.1 EEG offline preprocessing

EEG data were digitized with a sampling rate of 512 Hz. EEG data preprocessing was carried out using the EEGLAB toolbox (Delorme and Makeig, 2004; http://sccn.ucsd.edu/eeglab) in conjunction with FASTER. The data were initially bandpass filtered between 1 Hz and 95 Hz, notch filtered at 50 Hz and average referenced across all scalp electrodes. Resting data and data from the calibration and neurofeedback blocks were subsequently epoched into windows of 1000 ms. Data from the cSST were epoched from 500 ms prior to Go/Stop signal onset to 2000 ms after Go/Stop signal onset for Go trials and Stop trials, respectively. FASTER identified and removed artefactual (i.e., non-neural) independent components, removed epochs containing large artefacts (e.g., muscle twitches) and interpolated channels with poor signal quality. The remaining EEG data were then visually inspected by trained raters to ensure good quality and that any remaining noisy data were removed. Specifically, trained raters identified any remaining artefacts in independent components (e.g., eyeblinks) and epochs containing idiosyncratic muscle/movement or transient electrode artifacts, and interpolated any channels that were noisy throughout all epochs of a participant. After preprocessing, EEG data were transformed using the current source density method (CSD; https://psychophysiology.cpmc.columbia.edu/software/CSDtoolbox/index.html; Kayser and Tenke, 2006) which is a reference-free montage to attenuate the effect of volume conduction in scalp EEG.

#### 2.3.2 Time-frequency transformation

For all epochs, 2-dimensional representations of each electrode’s time-frequency were estimated using a complex Morlet wavelet (range of logarithmically spaced 4-10 cycles for 39 linearly spaced frequencies across 1-40 Hz). The squared magnitude of the convolved data was calculated to obtain power estimates. The power estimates were subsequently transformed to relative power. Power values of each given band from 1-28 Hz (Delta = 1-4 Hz, Theta = 5-7 Hz, Alpha = 8-12 Hz, Beta = 13-28 Hz) were expressed as a percentage of the total power within the spectrum (per channel and per given epoch). Beta bursts were extracted from non-relative time-frequency power estimates.

#### 2.3.3 Beta burst detection

Beta burst detection was performed according to the method described in Enz et al. (2021). For each time-frequency power matrix, local maxima were detected using the MATLAB function *imregionalmax*. Beta bursts were then defined as local maxima that exceeded a defined threshold of 2x median power of the entire time-frequency matrix (across all trials per participant). Time-frequency matrices were then divided into ~25.39 ms time bins (also for analysis of relative power). The first and last time bins were removed from all trials due to an edge artefact that can occur when applying the MATLAB function *imregionalmax* (it detects artefactual local maxima on the edges of the time-frequency matrix). Beta burst rate (the sum of the number of supra-threshold bursts) and Beta burst volume (the area under the curve of supra-threshold datapoints; see Enz et al., 2021) were then extracted per time bin. We also extracted the timing of the first Beta burst after the Stop/Go signal in the cSST EEG data.

#### 2.3.4 Selection of brain regions

For the statistical analysis, EEG data were averaged over clusters of four electrodes from different regions of the brain. To interrogate EEG data from the rIFC, data were averaged over the four individually selected electrodes over the right frontal scalp area that were used during the neurofeedback training. Further, we also averaged clusters of four electrodes over the left motor cortex (D19/C3, D20, D12, D11), the right motor cortex (B22/C4, B23, B31, B30) and the occipital cortex (A23/Oz, A24, A28, A27) to test the specificity or generalization of effects beyond the trained region.

### 2.4 Statistical analysis

#### 2.4.1 Behavioral analysis

Means and standard deviations were extracted for each participant for the following behavioral cSST measures: SSRT, intraindividual coefficient of variation (ICV), Go trial reaction time (RT), failed Stop trial RT, SSD, number of successful Stop trials, number of failed Stop trials, probability of successful stopping, probability of Go omissions, probability of choice errors. The SSRT was calculated using the integration method with replacement of Go omissions by the maximum RT (Verbruggen et al., 2019). All Go trials were included in the Go RT distribution, including Go trials with choice errors. Premature responses on failed Stop trials were included when calculating the probability of responding on a Stop trial and mean SSD. Participants with SSRT < 75 ms were excluded from all analyses. The ICV was calculated by dividing Go RT standard deviation by the mean Go RT.

For the 2-Back Task, the absolute number of target hits were calculated across both blocks (i.e., the maximum absolute number of target hits was 20).

#### 2.4.2 Neurofeedback training statistical analysis

For resting EEG and for the calibration and neurofeedback blocks, data were averaged over 37 x 25.39 ms time bins. For the cSST, 3 x 25.39 ms time bins were averaged to create ~75 ms time bins (6 time bins from −75 ms to 375 ms with respect to the Stop/Go signal). R (R Core Team, 2020) was used for all statistical analyses.

We first tested whether the neurofeedback training was effective. For this, we fit a linear mixed-effects model (LMM) using restricted maximum likelihood with relative Beta power over the rIFC as the outcome variable, with fixed effects of *Direction* (UP or DOWN) and *Timepoint* (Pre to Post neurofeedback training) and their two-way interaction, and with a random effect of *Participant*. We also calculated Cohen’s D for each effect. The models were fit for the Beta and Alpha groups separately. We then also conducted the same analysis with relative Alpha power as outcome variable. We also looked at the same outcome variables from the other three brain regions (left motor cortex, right motor cortex, occipital cortex). This analysis was repeated with Beta burst rate and Beta burst volume as outcome variables. We ran a post-hoc test for significant interactions using the *emmeans* function in R. All post-hoc tests are Bonferroni corrected at 0.05/2=0.025, correcting for the two directions (UP and DOWN).

Next, we looked at the effects of neurofeedback training on inhibitory control behavior. We again fit a LMM using restricted maximum likelihood with SSRT as the outcome variable, with fixed effects of *Direction* (UP or DOWN), *Timepoint* (Pre to Post neurofeedback training), *Rhythm* (Beta or Alpha) and their two- and three-way interactions, and with a random effect of *Participants*. We also calculated Cohen’s D for each effect.

We then interrogated the relationship between the magnitude of the Pre-Post change in SSRT and the extent to which relative Beta power was modulated during neurofeedback training. For this we fit a linear model by robust regression using an M-estimator, with change in SSRT being the dependent variable and change in relative Beta power being the independent variable. We repeated this analysis with N-Back score, Go RT and ICV as outcome variables.

Next, we looked at the effects of neurofeedback training on resting EEG data collected before and after. We fit the same three-way LMM with relative Beta power, Beta burst rate and Beta burst volume over the rIFC as outcome variables.

Last, we looked at the effect of neurofeedback training on the brain activity while engaging inhibitory control behavior. Again, the same three-way LMM was fit for the time bins around the average SSRT with relative Beta power, Beta burst rate, Beta burst volume and timing of first Beta burst over the rIFC as outcome variables.

### 2.5 Code and data accessibility

Custom written scripts and data summary files can be downloaded on the Open Science Framework at [URL to be inserted after acceptance].

## 3 Results

### 3.1 Spectral power in the Beta band over rIFC was modulated by Beta neurofeedback training

We first tested whether Beta and Alpha band spectral power was successfully modulated over rIFC by 6 days of neurofeedback in the trained directions (UP or DOWN), when quantified offline using optimal artefact rejection procedures. Linear mixed effects models were performed on EEG data recorded during BCI performance comparing spectral power at resting baseline on the first Day (Rest S1 Pre) to that at the end of the final (6^th^) Day (Block 6.4), within the two Beta subgroups. Beta power was significantly modulated from resting baseline on Day 1 to the final block of neurofeedback on Day 6, in a manner that differed depending on trained direction (**Figure 2A**). This was revealed by a Direction*Timepoint interaction (F[1,40.04]=5.98, p=0.019, d=0.77, n=44). N.B.: All following post-hoc tests are Bonferroni corrected at 0.05/2=0.025 and all means are shown as estimated marginal means (EMM) ± standard error. Beta power modestly increased for the UP group (Pre 62.0±3.03 %, Post 64.2±3.03 %; post-hoc test: t[39.1]=0.80, p=0.43) and significantly decreased for the DOWN group (Pre 59.5±3.65 %, Post 50.5±3.89 %; post-hoc test: t[41.5]=–2.47, p=0.018). The same pattern was evident when comparing data averaged within the calibration block performed immediately before training on Day 1 (Cal S1) to performance in the final block on Day 6 (F[1,38.18]=13.99, p=0.001, d=1.21, n=41; **Figure 2B**). Beta power modestly increased for the UP group (Pre 62.0±2.70 %, Post 64.2±2.67 %; post-hoc test: t[38.5]=0.99, p=0.33) and significantly decreased for the DOWN group (Pre 64.1±3.51 %, Post 52.3±3.51 %; post-hoc test: t[38.0]=–3.96, p=0.0003). The calibration block data was used to establish the midline of the on-screen display in the neurofeedback game, which participants were required to keep an avatar above (UP) or below (DOWN). In this block, participants rested for two minutes, then conducted a left-hand finger tapping movement for two minutes in order to establish the full range of raw values associated with synchronization and desynchronization of the individual participant’s Beta rhythm. **Figure 2A-B** show the time course of Beta power for S1 and S6 compared to resting baseline (**Figure 2A**) and calibration baseline (**Figure 2B**). See Supplementary Results 1 for acute within session modulation for Day 1 and Day 6.

**Figure 2:**
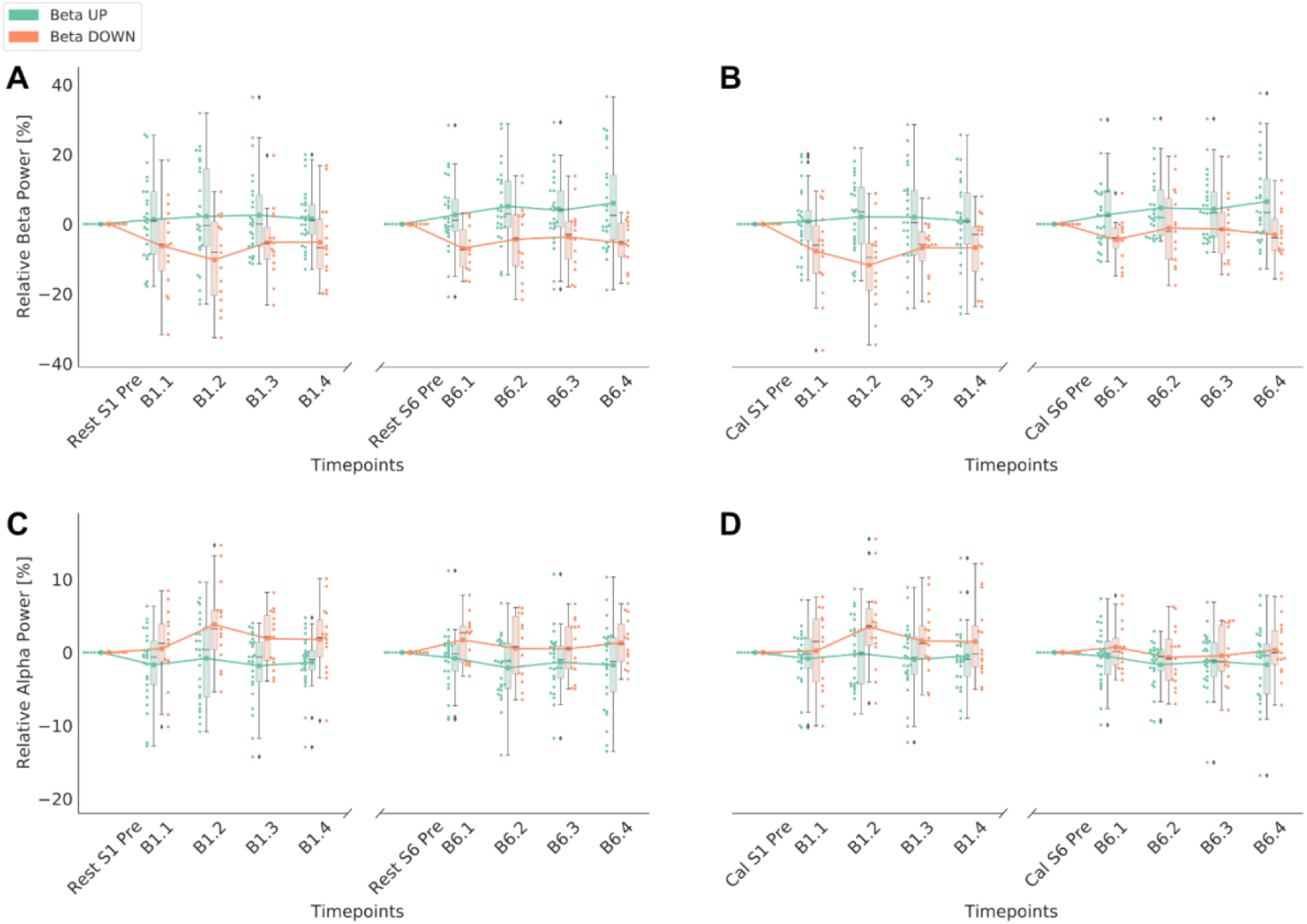
Neurofeedback training performance of Beta groups. Performance of first and last session blocks is shown for Beta UP and Beta DOWN groups separately. Time course of relative power is corrected to the respective baseline. The boxplots show the medians and quartiles of the data, the whiskers extend to the rest of the distribution, except for points that are determined to be outliers. The swarm plots show individual datapoints and the line plots connect the means of each block. **A)** Relative Beta power is shown relative to the resting baseline before the first training block on Day 1. **B)** Relative Beta power is shown relative to the calibration baseline before the first training block on Day 1. **C)** Relative Alpha power is shown relative to the resting baseline before the first training block on Day 1. **D)** Relative Alpha power is shown relative to the calibration baseline before the first training block on Day 1.

Spectral power in the Beta band over rIFC was not significantly modulated from resting baseline during Alpha BCI training (Direction*Timepoint interaction: F[1,31.94]=1.42, p=0.24, d=0.42, n=37; **Figure 3C**). When comparing end of training (B6.4) to the calibration block on Day 1, the Direction*Timepoint interaction was significant (F[1,33.10]=4.38, p=0.044, d=0.73, n=36) but post-hoc tests revealed that neither the UP nor DOWN groups showed significant modulation of the Beta rhythm from the calibration baseline (all p>0.13; **Figure 3D**). Power in the Alpha band was similarly not modulated during Alpha neurofeedback (all Direction*Timepoint interactions p>0.42; **Figure 3A-B**), suggesting that training Alpha over rIFC was not achieved using this protocol.

**Figure 3:**
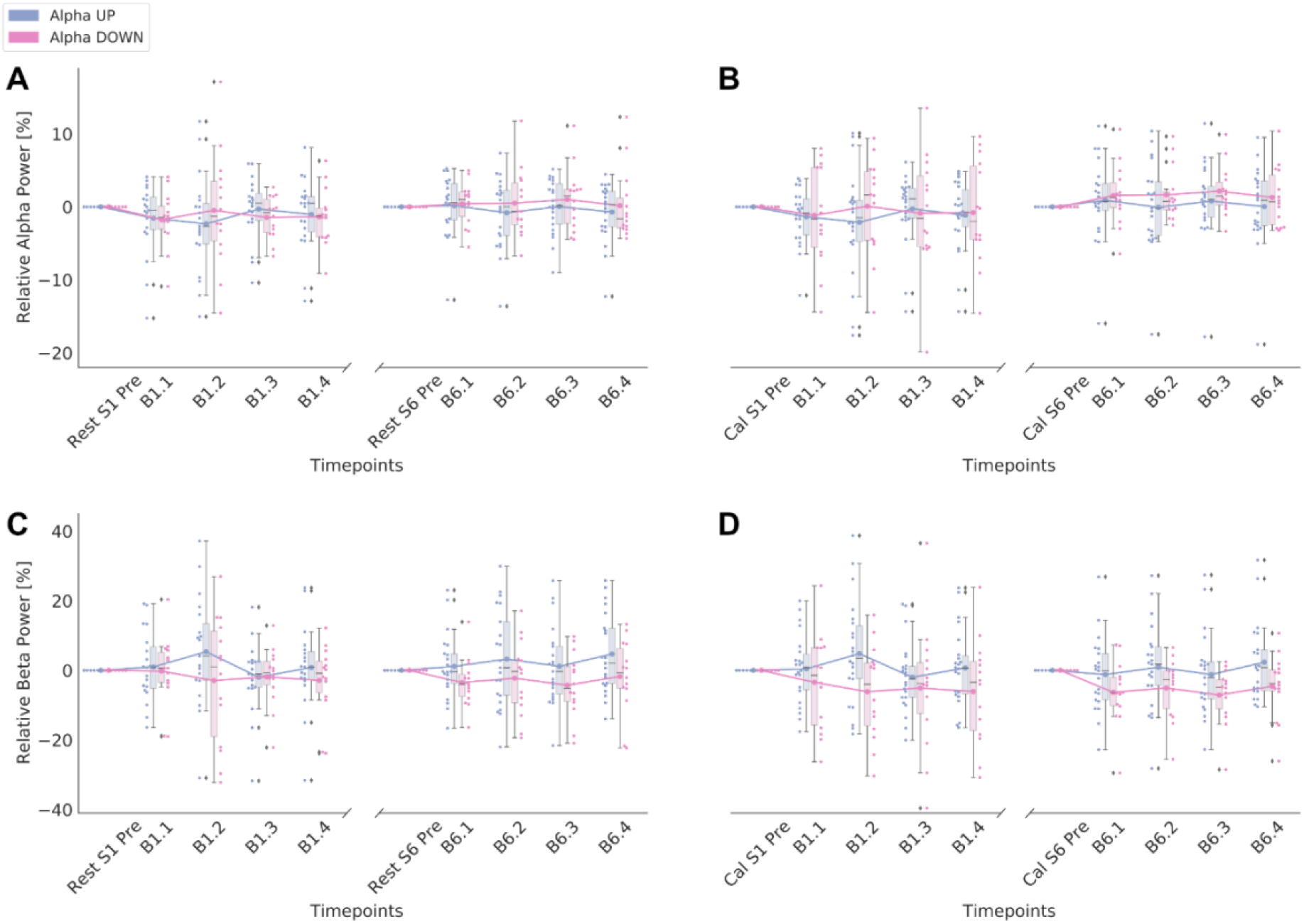
Neurofeedback training performance of Alpha groups. Performance of first and last session blocks is shown for Alpha UP and Alpha DOWN groups separately. Time course of relative power is corrected to the respective baseline. The boxplots show the medians and quartiles of the data, the whiskers extend to the rest of the distribution, except for points that are determined to be outliers. The swarm plots show individual datapoints and the line plots connect the means of each block. **A)** Relative Alpha power is shown relative to the resting baseline before the first training block on Day 1. **B)** Relative Alpha power is shown relative to the calibration baseline before the first training block on Day 1. **C)** Relative Beta power is shown relative to the resting baseline before the first training block on Day 1. **D)** Relative Beta power is shown relative to the calibration baseline before the first training block on Day 1.

### 3.2 Cross frequency effects and modulation of neural oscillations beyond rIFC

We first investigated whether power in the Alpha band was modulated during neurofeedback targeting the Beta rhythm (**Figure 2C-D**). When comparing calibration baseline Alpha power on Day 1 to Alpha power during neurofeedback attempting to regulate Beta, significant modulation was detected at the end of Day 6 (F[1,38.33]=4.78, p=0.035, d=-0.71, n=41). Alpha power modestly decreased for the UP group (Pre 16.1±1.26 %, Post 15.8±1.24 %; post-hoc test: t[38.5]=-0.21, p=0.83) and significantly increased for the DOWN group (Pre 17.0±1.64 %, Post 20.5±1.64 %; post-hoc test: t[38.0]=2.60, p=0.013). It is notable however that although this demonstrates that Alpha was modulated during training based upon neurofeedback of the Beta rhythm, the direction of change was opposite (Alpha decreased in the Beta UP training and vice versa). Also, the absolute effect sizes for Alpha modulation during Beta training range from 0.23-0.71, whereas Beta modulation absolute effect sizes were substantially larger (0.61-1.21). On the final Day of training, Alpha power was not modulated acutely (i.e., within session) during Beta training when comparing power during the final neurofeedback block to the resting baseline on the same day (F[1,38.75]=3.16, p=0.08, d=-0.57, n=41), nor to the calibration block (F[1,37.06]=1.35, p=0.25, d=-0.38, n=41). Thus, the effects of Beta training were largely selective to the Beta rhythm.

To investigate effects spanning beyond the trained cluster of electrodes over rIFC, we tested whether Beta Power at three other scalp sites was modulated during neurofeedback of Beta signals recorded from rIFC. We chose clusters of four electrodes over right and left motor cortex and occipital cortex for comparison and performed mixed effects models testing for Direction*Timepoint interactions, as before. No significant interactions in any of the three regions were detected when comparing resting baseline Beta power to Beta power during the final neurofeedback block on Day 6 (all p>0.12). However, when comparing Beta power from the initial calibration block on Day 1 to power during the final block on Day 6, significant Direction*Timepoint interactions were revealed for both right (F[1,38.45]=8.46, p=0.006, d=0.94, n=41) and left (F[1,38.36]=6.58, p=0.014, d=0.83, n=41) motor regions, but no modulation was evident in the occipital region (F[1,35.03]=0.29, p=0.59, d=-0.18, n=41). For the right motor region, Beta power modestly increased for the UP group (Pre 56.9±2.34 %, Post 58.9±2.31 %; post-hoc test: t[38.5]=0.92, p=0.36) and significantly decreased for the DOWN group (Pre 59.4±3.04 %, Post 51.3±3.04 %; post-hoc test: t[38.0]=–2.96, p=0.005). For the left motor region, Beta power modestly increased for the UP group (Pre 57.6±2.29 %, Post 59.5±2.27 %; post-hoc test: t[38.4]=1.13, p=0.26) and significantly decreased for the DOWN group (Pre 57.4±2.99%, Post 52.3±2.99 %; post-hoc test: t[38.0]=–2.37, p=0.023). Topoplots for post-training minus pre-training are shown for each training group for both Beta power (**Figure 4**) and Alpha power (**Figure S1**).

**Figure 4:**
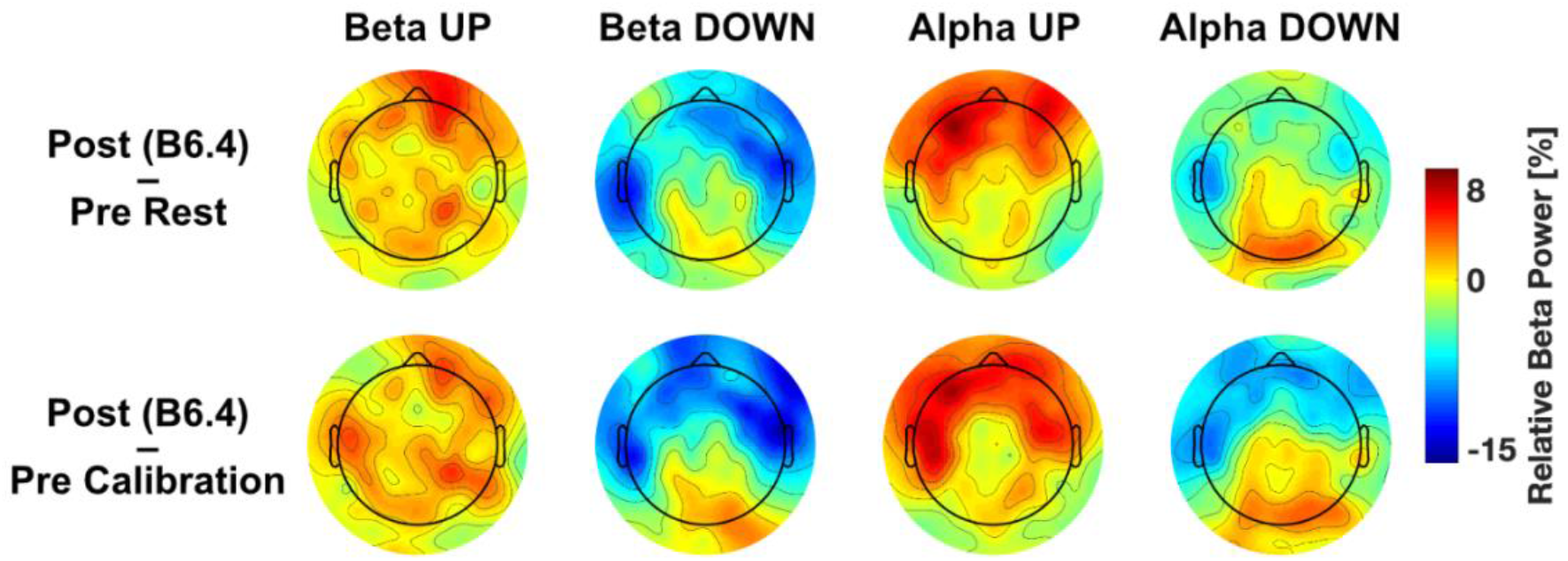
Change in Beta power pre-to post-neurofeedback training. Topoplots show relative Beta power for the last block of neurofeedback training (Post B6.4) minus resting EEG before first block of neurofeedback training (Pre Rest) as well as for the last block of neurofeedback training (Post B6.4) minus calibration block (Pre Calibration). Topoplots are shown separately for each group (Beta UP, Beta DOWN, Alpha UP, Alpha DOWN).

### 3.3 Change in inhibitory control behavior modulated by neurofeedback training

Behavioral data of the cSST are displayed in **Table S1**. To assess whether neurofeedback training had any effect upon inhibitory control behavior in the cSST (i.e., upon SSRT, the speed of inhibitory control), we performed a mixed effects model with three fixed effects; Rhythm (Alpha or Beta), Direction (UP or DOWN) and Timepoint (pre or post training). There was no 3-way Rhythm*Direction*Timepoint interaction (F[1,54.88]=2.32, p=0.13, d=0.41, n=71). There was a fixed effect of Timepoint (F[1,54.88]=4.35, p=0.042, d=0.28, n=71), revealing that SSRTs generally improved over time regardless of BCI training type (EMMs averaged over levels of Rhythm and Direction: Pre 166±7.86 ms, Post 147±8.54 ms). **Figure 5** shows the mean pre and post SSRTs for each group.

**Figure 5:**
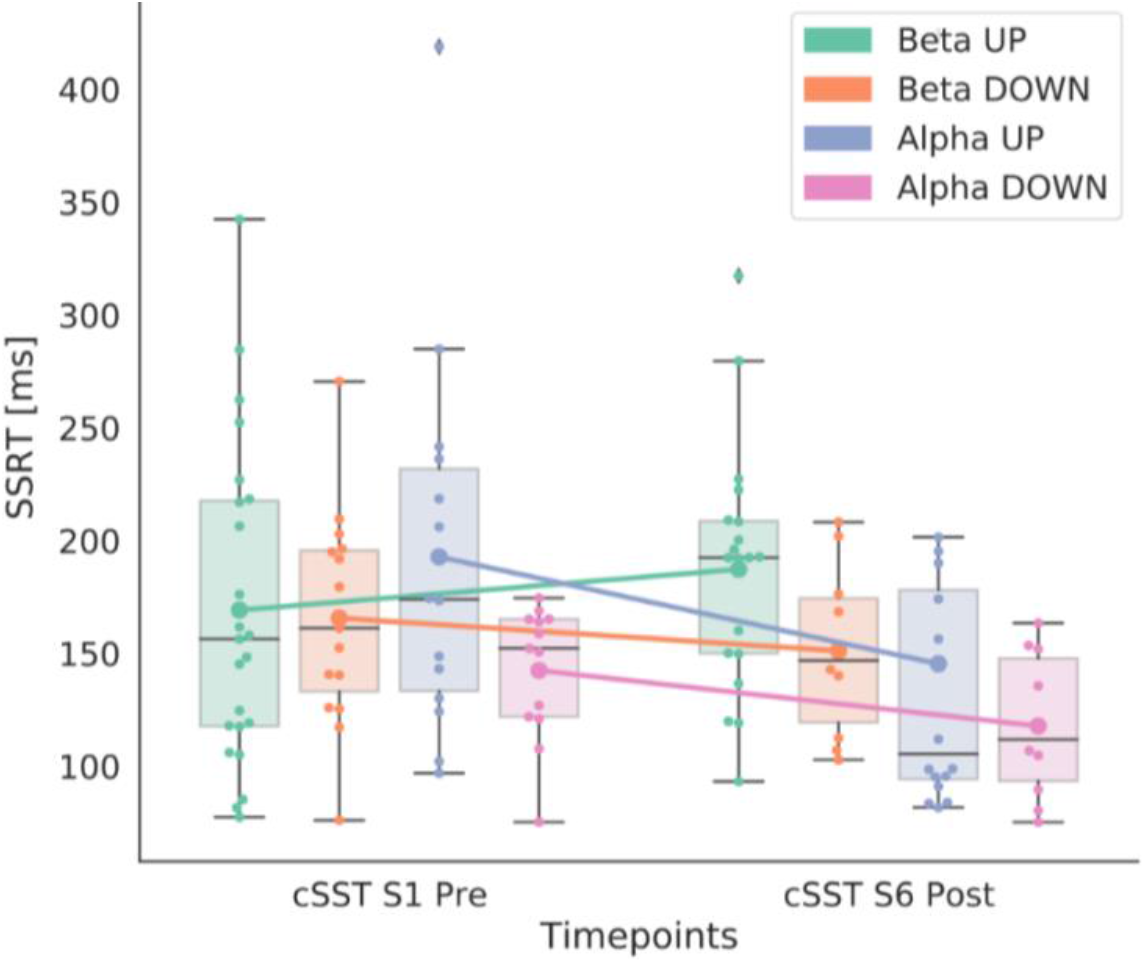
SSRTs of each training group. The SSRTs are shown for each training group for pre (Day 1) and post (Day 6) neurofeedback training. The boxplots show the median and quartiles of the data, the whiskers extend to the rest of the distribution, except for points that are determined to be outliers. The swarm plots show individual datapoints and the line plots connect the means of each block. Abbreviations: SSRT: Stop signal reaction time; cSST: conditional stop signal task.

To investigate whether each individual’s pre-post change in SSRT could be predicted by the extent to which their Beta rhythm was modulated, we performed Robust Regression analyses with change in SSRT as outcome variable and change in Beta rhythm (resting baseline from Day 1 to final block on Day 6) as predictor. The extent of change in the Beta rhythm as a result of training did not significantly predict improvement in SSRT from Pre-Post (slope=0.81, df=25, F=2.06, p=0.16, n=22; **Figure 6A**). The same was evident when tested at the right motor electrode cluster (slope=1.15, df=25, F=2.76, p=0.11, n=22; **Figure 6B**), left motor cluster (slope=1.22, df=25, F=2.38, p=0.14, n=22; **Figure 6C**), and occipital cluster (slope=0.46, df=25, F=0.41, p=0.53, n=22; **Figure 6D**). Training related change in Alpha power for those training Alpha rhythms was not predictive of behavioral change in SSRT (all p>0.26; **Figure 6E-H**).

**Figure 6:**
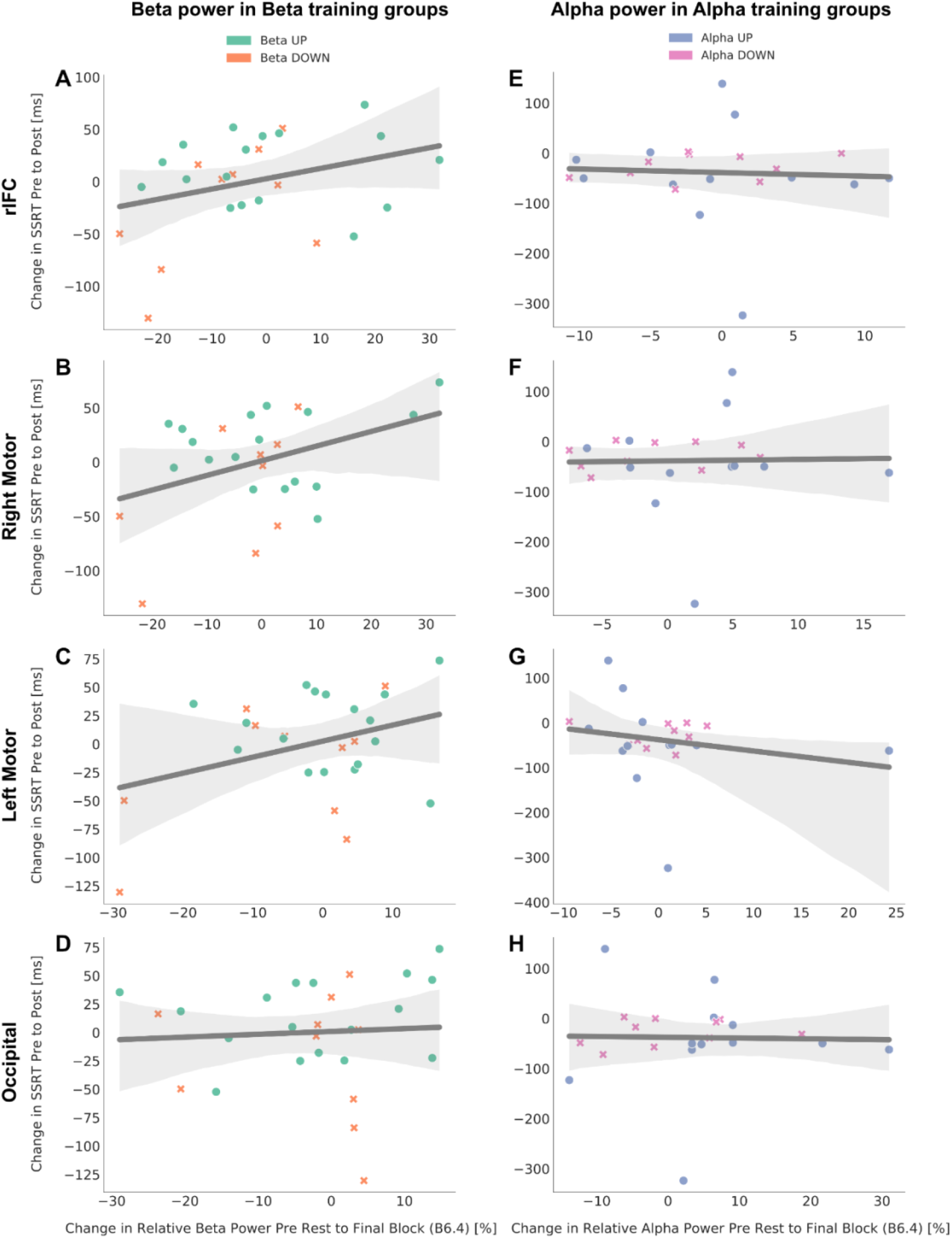
Association between change in SSRT and change in relative power. Each plot shows the linear regression fit line of the data in dark grey as well as the confidence interval in light grey. Please note the difference in scale across all plots. The associations are shown for an average of four electrodes in four different brain regions (rIFC, right motor, left motor, occipital). **A-D)** show the association between change in SSRT and change in relative Beta power in the Beta groups (UP and DOWN). **E-H)** show the association between change in SSRT and change in relative Alpha power in the Alpha groups (UP and DOWN). Abbreviations: SSRT: stop signal reaction time; rIFC: right inferior frontal cortex.

We additionally tested whether neurofeedback training impacted other aspects of cognitive function, including working memory (2-Back Task), processing speed (Go RT; Go reaction times in the cSST) and performance variability (ICV; intra-coefficient of variation in the cSST). No 3-way interactions emerged between Rhythm*Direction*Timepoint (all p>0.41), but for Go RT there was a significant 2-way Rhythm*Timepoint interaction (F[1,75.34]=4.82, p=0.031, d=0.47, n=82). Post-hoc tests are indicating a general improvement in speed predominantly in the Beta group, revealing that Go RT significantly decreased for the Beta group (Pre 484±7.07 ms, Post 463±7.31 ms; post-hoc test: t[5.40]=3.44, p=0.001) but did not significantly decrease for the Alpha group (Pre 480±7.65 ms, Post 478±7.79 ms; post-hoc test: t[74.4]=0.24, p=0.82).

### 3.4 Resting spectral Beta power was not altered by neurofeedback training

Comparing resting EEG data from before training (Rest S1 Pre), after Day 1 of training (Rest S1 Post) and Post training on Day 6 (Rest S6 Post), mixed effects models revealed no significant Direction*Timepoint interactions for the Beta group (F[2,79.42]=0.34, p=0.71, n=44), suggesting that training related modulations of Beta power were only evident when engaged in the task but did not lead to a lasting change in the background (resting) tonic Beta level.

### 3.5 Modulation of Beta burst characteristics by training the tonic Beta Rhythm

We investigated whether training to modulate tonic Beta Power over rIFC has consequences for Beta burst characteristics. Beta burst rate was not altered during or after training at any of the timepoints tested (all p>0.277). Burst volume was significantly modulated at the end of the first Day of training when comparing burst volume in the last block of Day 1 to that detected in the calibration block on the same Day (Direction*Timepoint interaction: F[1,37.48]=6.14, p=0.02, d=0.81, n=40). Burst volume significantly increased for the UP group (Pre 5962±1744 a.u., Post 9769±1771 a.u.; post-hoc test: t[36.6]=2.20, p=0.03) and modestly decreased for the DOWN group (Pre 8560±2311 a.u., Post 5336±2251 a.u.; post-hoc test: t[37.0]=–1.43, p=0.16).

### 3.6 Brain activity while performing the cSST was not modified following training

There were no significant differences in neural activity (Beta power, burst rate, burst volume, timing of first burst) recorded during cSST performance at the start of the first day of training compared to the end of Day 6 of training (all p>0.40).

## 4 Discussion

We have demonstrated here that using neurofeedback in a BCI it is possible to train human participants to self-regulate their Beta rhythm over the rIFC, but that this has no observable consequences upon subsequent inhibitory control behavior. Participants trained over a 6-day period to upregulate or downregulate the amplitude of their Beta rhythm over rIFC, resulting in the predicted directional changes to Beta power. Concomitant changes at other (untrained) scalp regions and other frequency bands were of lower magnitude, indicating good specificity of the neurofeedback protocol for modulating the trained rhythm, direction and region. The extent to which each individual’s SSRT changed pre-post training was however, not predicted by the magnitude of their training-related change in Beta over rIFC. This was also not the case for the control group undergoing Alpha training. Although the right frontal Beta rhythm has been repeatedly implicated as a key component of the brain’s inhibitory control system, the present data suggest that improving the ability to self-regulate the rhythm does not result in behavioral change in an inhibitory control task.

Training related modulation of the Beta rhythm was only manifest during the neurofeedback task and did not alter EEG signals measured subsequently at rest or during cSST task performance. The current experimental design did not permit us to investigate whether online (i.e., during cSST task performance) self-regulation of the Beta rhythm would impact upon behavior, although this may be an interesting future extension of the work. Additionally, in the BCI task, neurofeedback was provided on the amplitude of the tonic (background) Beta rhythm. Further analyses revealed that this style of regulation of tonic Beta power had no impact on the rate or volume of transient burst-like high amplitude events in the Beta frequency range. This adds weight to the emerging view that so called ‘Beta bursts’ are a phenomena distinct from the ongoing background or ‘tonic’ oscillation at the same frequency (Bonaiuto et al., 2021; Little et al., 2019). Timing and magnitude of Beta bursts critically impact upon subsequent motor performance (Little et al., 2019) and whether attempts to inhibit a response are successful or not (Enz et al., 2021; Wessel, 2020). Although participants learned to modulate the amplitude of their Beta rhythm over rIFC, this gradual, tonic background change in Beta during the distinct 3-minute neurofeedback blocks had no impact upon subsequent inhibitory control behavior. In order to modify Beta bursts using the BCI, it may be necessary to provide feedback specifically tailored to detect and influence bursting in real-time (online), rather than simply of generalized (and offline) regulation of tonic Beta (e.g., He, 2020).

In the protocol used in the current study, neurofeedback targeting downregulation of Beta oscillations was more impactful than upregulation. It is likely that downregulation is simply easier for participants to perform, as it is known that engaging a brain region in a mental process (such as motor imagery), tends to lead Beta (and Alpha) to desynchronize in the region (Jensen & Mazaheri, 2010). Over the 6-day training period, participants learned to tailor their mental imagery strategies to optimally engage rIFC in order to achieve tangible real-time control over the movement of the avatar on screen. Our results leave open the possibility that it may be the ability to flexibly engage (and remove) Beta oscillations in the form of precisely timed bursts that predicts behavioral performance, rather than the tonic level per se.

While the neurofeedback training we employed was effective for regulating the Beta rhythm over rIFC, Alpha modulation at this scalp location was not achieved. The lack of Alpha modulation over rIFC may be due to the fact that Beta is the predominant resonating frequency in this location, and has been repeatedly implicated in the functioning of this region (Schaum et al., 2021; Sundby et al., 2021; Swann et al., 2009; Swann et al., 2012; Wagner et al., 2017). Further, for all participants (even those in the Alpha group) we performed the same functional localizer to detect the precise cluster of electrodes corresponding to the right frontal scalp location showing most substantial Beta synchronization during the cSST. This cluster of electrodes, selected for exhibiting strong Beta activity during inhibitory control, was used to tailor the BCI neurofeedback for both Alpha and Beta groups. Optimizing the BCI for Beta using this method may further explain why Alpha modulation was not achieved at the rIFC site.

Previous studies using implanted electrodes have reported that positive effects upon motor behavior could be achieved in macaques (Khanna & Carmena, 2017) and humans with Parkinson’s disease (Bichsel et al., 2021) using BCI to train self-regulation of the brain’s Beta rhythm. Here we build upon and extend these initial findings by making the advance to non-invasive scalp recorded EEG signals in humans, demonstrating that volitional modulation of Beta oscillations was achieved within 6 days of training. The BCI neurofeedback protocol demonstrated good spatial and temporal specificity, modulating primarily the targeted region, rhythm and direction. The lack of behavioral consequences further adds weight to the emerging picture in recent research showing that the right frontal Beta signature associated with stopping may not exert a direct functional influence upon the behavior. Indeed, Errington et al. (2020) demonstrated using depth electrodes in macaques that while Beta bursts were associated with inhibitory control, successful stopping could occur even on trials where no bursts were detected. They also highlighted that the occurrence of Beta bursts during Stop trials was generally very low (~15% of trials), and as such may only represent one component of a more complex neural mechanism underlying inhibitory control.

Using non-invasive BCI technology, volitional and causal self-regulation was achieved without the need for exogenous stimulation, paving the way for easier real-world application of neuromodulation to alter brain rhythms experimentally. The present data suggest, however, that offline neurofeedback training of the tonic Beta rhythm may not serve as a useful therapeutic target for disorders with dysfunctional inhibitory control as their basis.

## Acknowledgements

The authors would like to thank Emma Wall, Aisling Martin and David M. Cole for assistance during data collection. Nadja Enz is supported by Irish Research Council postgraduate scholarship GOIPG/2018/537. Robert Whelan was supported by Science Foundation Ireland (16/ERCD/3797); European Foundation for Alcohol Research (ERAB); Brain & Behavior Research Foundation (23599); Health Research Board HRAPOR-2015-1075. Kathy L. Ruddy would like to acknowledge funding from the Irish Research Council GOIPD/2017/798 and Health Research Board, Ireland HRB-EIA-2019-003.

