## Supplementary Material for "Self-regulation of the brain’s right frontal Beta rhythm using a brain-computer interface"

**Short title:** Self-regulation of the right frontal Beta rhythm

**Author names:** Nadja Enz[1], Jemima Schmidt[1], Kate Nolan[1], Matthew Mitchell[1], Sandra Alvarez Gomez[1], Miryam Alkayyali[1], Pierce Cambay[1], Magdalena Gippert[1], Robert Whelan*[1,2], Kathy L. Ruddy*[1]

***Equal contribution***

**Affiliations:** [1]School of Psychology and Institute of Neuroscience, Trinity College Dublin, Dublin, Ireland, [2]Global Brain Health Institute, Trinity College Dublin, Dublin, Ireland

**Corresponding Authors:**

Kathy L. Ruddy**,** Lloyd Institute, Trinity College Institute of Neuroscience and School of Psychology, Trinity College Dublin, College Green, Dublin 2, Ireland

Robert Whelan, Lloyd Institute, Trinity College Institute of Neuroscience and School of Psychology, Trinity College Dublin, College Green, Dublin 2, Ireland

**Acknowledgements**: The authors would like to thank Emma Wall, Aisling Martin and David M. Cole for assistance during data collection. Nadja Enz is supported by Irish Research Council postgraduate scholarship GOIPG/2018/537. Robert Whelan was supported by Science Foundation Ireland (16/ERCD/3797); European Foundation for Alcohol Research (ERAB); Brain & Behavior Research Foundation (23599); Health Research Board HRAPOR-2015-1075. Kathy L. Ruddy would like to acknowledge funding from the Irish Research Council GOIPD/2017/798 and Health Research Board, Ireland HRB-EIA-2019-003.

**Competing interests:** The authors declare no competing financial or non-financial interests.

### 1 Supplementary Results

**Supplementary Results 1: Acute modulation of Beta power on first and last Day of Beta training**

There was no acute (within session) modulation of Beta power in the first training block on the first day when comparing against the resting baseline (F[1,38.66]=2.58, p=0.12, d=0.52, n=44), but there was significant acute modulation when comparing against the calibration baseline (F[1,38.0]=5.65, p=0.002, d=0.77, n=40), revealing that Beta power modestly increased for the UP group (EMM (Estimated marginal mean) ± SE: Pre 61.6±2.87 %, Post 62.4±2.87 %; post-hoc test: t(38.0)=0.36, p=0.72) and significantly decreased for the DOWN group (Pre 64.1±3.71 %, Post 56.3±3.71 %; post-hoc test: t(38)=-2.73, p=0.01). On the final Day of training, significant acute modulation of Beta was evident, comparing Beta power in the final Block to resting baseline at the beginning of the same session (Rest S6 Pre; F[1,38.24]=8.97, p=0.005, d=0.97, n=41), revealing that Beta power significantly increased for the UP group (Pre 58.2±2.81 %, Post 64.2±2.81%; post-hoc test: t(38.0)=2.54, p=0.015) and modestly decreased for the DOWN group (Pre 58.2±3.77 %, Post 52.3±3.69 %; post-hoc test: t(38.8)=-1.85, p=0.072). Significant acute modulation on the final Day was also apparent when comparing to the calibration block data on the same Day (Cal S6; F[1,37.50]=6.07, p=0.019, d=0.81, n=41), revealing that Beta power significantly increased for the UP group (Pre 58.1±2.80 %, Post 64.2±2.76 %; post-hoc test: t(38.4)=2.69, p=0.011) and modestly decreased for the DOWN group (Pre 55.4±3.64 %, Post 52.3±3.64 %; post-hoc test: t(38.0)=-1.04, p=0.31).

### 2 Supplementary Figures

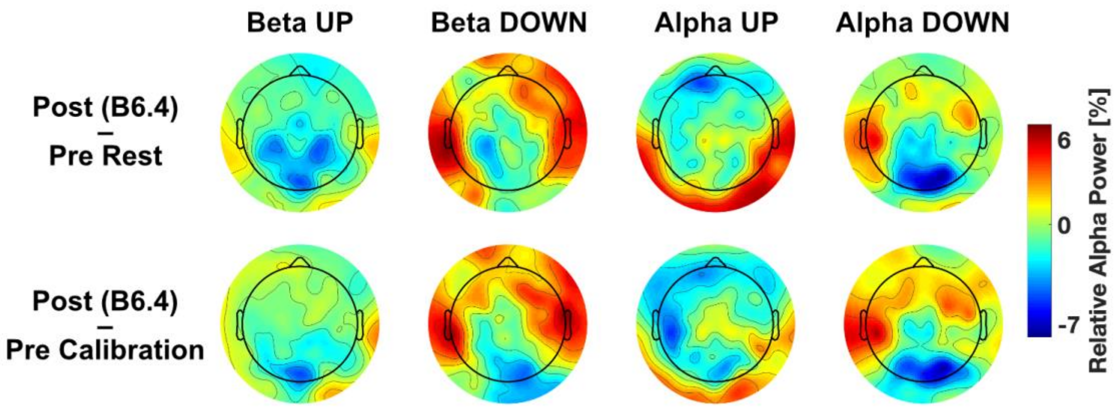


**Figure S1: Change in Alpha power pre- to post-neurofeedback training.** Topoplots show relative Alpha power for the last block of neurofeedback training (Post B6.4) minus resting EEG before first block of neurofeedback training (Pre Rest) as well as for the last block of neurofeedback training (Post B6.4) minus calibration block (Pre Calibration). Topoplots are shown separately for each group (Beta UP, Beta DOWN, Alpha UP, Alpha DOWN).

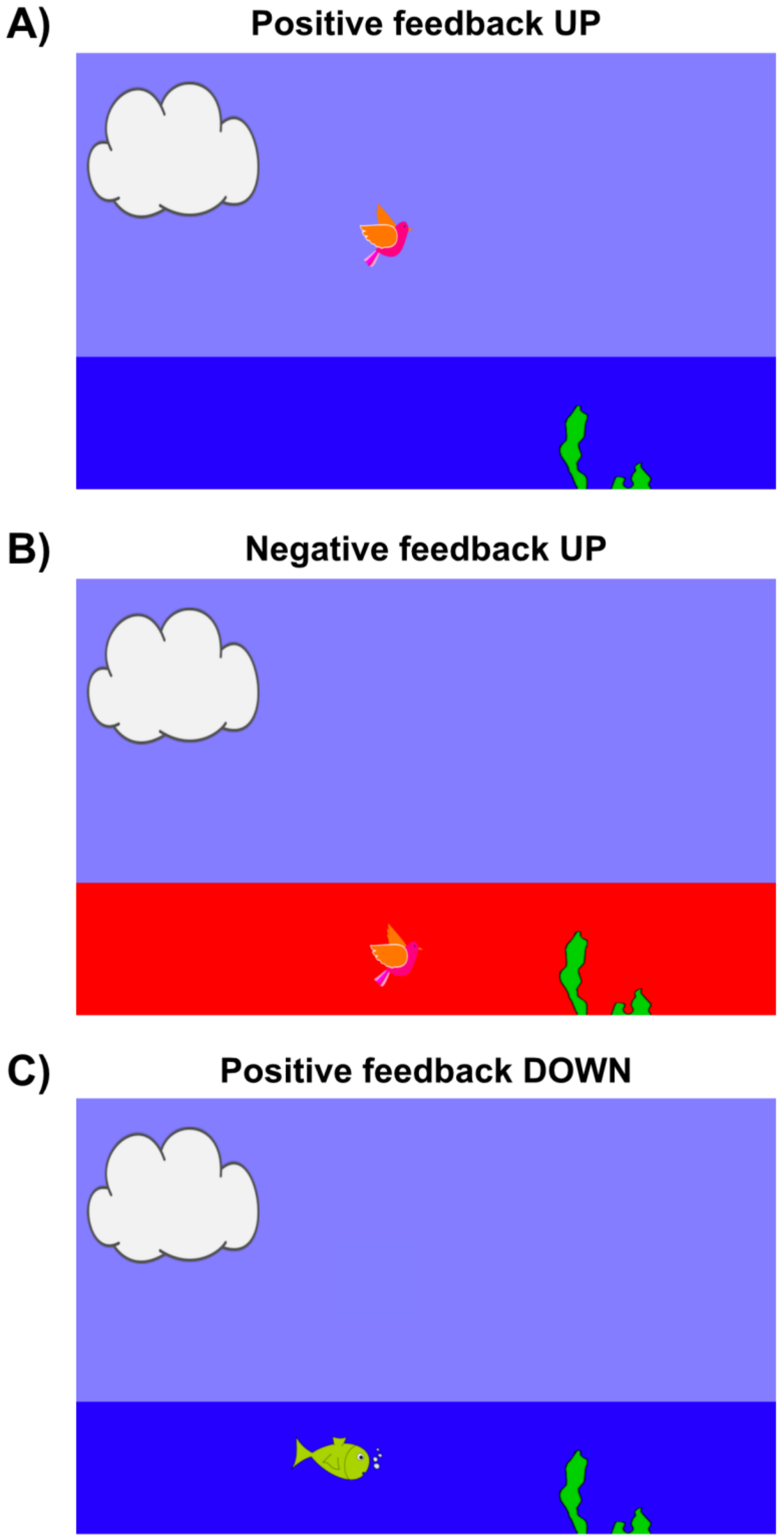


**Figure S2: Visual neurofeedback.** A) Positive feedback for the UP groups. The participant's real-time power estimate output is visualized in form of a bird. If the bird is in the sky, the background stays normal. B) Negative feedback for the UP groups. If the bird reaches the sea, the background turns red to give negative feedback. C) Positive feedback for the DOWN groups. The participant's real-time power estimate output is visualized in form of a fish.

### 3 Supplementary Table

**Table S1. Behavioural data of the cSST pre, acute and post neurofeedback training.** Means and standard deviations are reported. Abbreviations: cSST: conditional stop signal task; SSRT: stop signal reaction time; SSD: stop-signal delay; RT: reaction time.

| | Pre SST | Acute SST | Post SST |
|---|---|---|---|
| SSRT (ms) | 168 (65) | 167 (77) | 156 (64) |
| SSD (ms) | 301 (60) | 288 (71) | 307 (72) |
| Go RT (ms) | 482 (43) | 465 (44) | 468 (49) |
| Failed stop RT (ms) | 466 (52) | 458 (53) | 463 (61) |
| Successful stop (%) | 59.5 (10.5) | 55.5 (11.6) | 59.1 (12.5) |
| Go omission (%) | 5.4 (8.1) | 3.5 (5.5) | 2.3 (4.7) |
| Choice errors (%) | 1.1 (1.1) | 1.0 (1.4) | 0.9 (1.1) |

*bioRxiv versions accessed prior to 22-Nov-2021 contained a data error*
